# The Role of Conjunctive Representations in Prioritizing and Selecting Planned Actions

**DOI:** 10.1101/2022.05.09.491164

**Authors:** Atsushi Kikumoto, Ulrich Mayr, David Badre

## Abstract

For flexible goal-directed behavior, prioritizing and selecting a specific action among multiple candidates is often important. Working memory has long been assumed to play a role in prioritization and planning, while bridging cross-temporal contingencies during action selection. However, studies of working memory have mostly focused on memory for single components of an action plan, such as a rule or a stimulus, rather than management of all of these elements during planning. Therefore, it is not known how post-encoding prioritization and selection operate on the entire profile of representations for prospective actions. Here, we assessed how such control processes unfold over action representations, highlighting the role of conjunctive representations that nonlinearly integrate task-relevant features during maintenance and prioritization of action plans. For each trial, participants prepared two independent rule-based actions simultaneously, then they were retro-cued to select one as their response. Prior to the start of the trial, one rule-based action was randomly assigned to be high priority by cueing that it was more likely to be tested. We found that both full action plans were maintained as conjunctive representations during action preparation, regardless of priority. However, during output selection, the conjunctive representation of the high priority action plan was more enhanced and readily selected as an output. Further, the strength of conjunctive representation was related to behavioral interference when the low priority action was tested. Thus, multiple integrated representations were maintained for upcoming actions and served as the target of post-encoding attentional selection mechanisms to prioritize and select an action from those in working memory.

## Introduction

Guiding actions in a complex environment often involves specification and selection of multiple candidate plans (Cisek 2007). Our ability to prepare and then single-out a planned action allows us to keep actions for several potential scenarios ready, and then decide which to execute based on their adaptive utility in the current context. Such flexible action control requires separate mechanisms for rapidly translating incoming sensory information into action-oriented plans in a context-dependent manner and for deciding to enact those plans as behavioral output (Cisek and Kalaska 2005; Gallivan et al. 2015). To this end, working memory is thought of as a core interface, both for assembling and maintaining task-relevant plans, as well as gating a subset of this information to use when actions become relevant (Chatham and Badre 2015; Myers et al. 2017; Kriete and Noelle 2011; Frank and Badre 2012; Badre and Frank 2012).

Top-down prioritization and selection of the encoded information has been commonly recognized as a function of working memory (Souza and Oberauer 2016; Gazzaley and Nobre 2012). Retrospectively directing attention to specific memories (e.g., via retro-cues) makes the targeted content more accessible for later retrieval and more resilient against interference, while increasing its neural decodability (LaRocque et al. 2013; Yu et al. 2020; Ester et al. 2018). However, although the primary function of working memory is to guide future actions rather than passively store past sensations (Miller et al. 2018; Olivers and Roelfsema 2020; Nobre and Stokes 2019), control over working memory has been studied mostly with regard to sensory representations (e.g., words, letters, symbols, faces, scenes, and so forth) that are independent from their prospective goals and actions (Myers et al. 2017; Heuer et al. 2020; van Ede 2020). Because realistic goal-directed actions are supported by multiple task features, which often influence to one another dynamically (e.g., a hierarchical gating; see Rac-Lubashevsky and Frank 2021; Ranti et al., 2016; Frank and Badre, 2012), it remains an open question how the full complement of task-relevant features comprising an action plan are managed together within working memory.

One important representational format that may support control over planned actions is a conjunctive representation that evolves over the course of action planning (Hommel 2019; Frings et al. 2020). This conjunctive representation, also termed an event or task file, binds critical task-relevant features **—**including not only the sensory representation (i.e., stimulus), but also the action rule (i.e., context) and response**—**into an integrated representation that uniquely couples sensory and motor information for a specific goal (Mayr and Bryck 2005). Recent studies that applied a decoding analysis to the distributed pattern of EEG activity revealed that humans form such conjunctive representations while preparing actions. Furthermore, trial-to-trial fluctuation in the strength of a conjunction strongly correlated with efficient action control (e.g., stopping of actions) over and above other task-relevant representations (Kikumoto and Mayr 2020; Kikumoto 2021; Rangel, Hazeltine, and Wessel, 2022).

Because a conjunctive representation uniquely specifies the to-be-executed action, it may be an efficient, output-oriented format for maintenance, prioritization and retrospective selection of the planned actions for future behavior (Badre et al. 2021). Yet, previous studies have focused on a single episode of action selection. Thus, it is unknown whether more than one competing conjunctive representation can be maintained in parallel. If multiple conjunctive representations could be prepared concurrently, action prioritization may primarily modulate output gating of prepared of action plans. In contrast, if there is a capacity limit on holding multiple event-file like representations, maintenance of a given action plan should be compromised by trying to simultaneously prepare for alternative action plans.

In the current study, we investigated the role of conjunctive representations in the maintenance and prioritization of multiple candidate action plans for an upcoming response. Participants prepared two independent actions with unique stimulus-response mappings and chose one of them as a final response based on a retrospective cue. One of the actions was, on average, more likely to be tested, encouraging selective prioritization of that action until output selection.

We hypothesized (a) that future actions are maintained concurrently in the form of multiple conjunctive representations during action preparation and (b) that conjunctions are the primary target of prioritization and selection once a planned action is selected (over and above individual components of the action plan like the stimulus, rule, or response). To preview our results, RTs and errors indicated that participants prepared both actions but held the higher priority action in a privileged state. By applying a time-resolved representational similarity analysis (RSA) to EEG, we found that conjunctive representations of both high and low priority actions were maintained up until the moment of selection. Further, the conjunctive representation of the high priority action was more enhanced and readily selected as an output, and it produced interference when the low priority action was selected instead. Such a modulatory effect by priority was weak for stimulus representations, suggesting that the conjunction as a primary target of prioritization and selection. Thus, consistent with our hypothesis, multiple candidate actions are maintained, prioritized, and selected as integrated, but independent action representations.

## Materials and Methods

### Participants

Twenty-six participants (16 females; mean age: 23.2 years) were recruited from the University of Oregon student body. All participants had normal or corrected-to-normal vision, and had no history of neurological or psychiatric disorders. They were compensated $10 per hour plus additional performance-based incentives (see *Behavioral Procedure* section). Participants underwent informed consent using procedures approved by the Human Subjects Committee at the University of Oregon. After preprocessing the EEG data, two participants were removed and were not analyzed further due to excessive amounts of artifacts (i.e., more than 25% of trials; see *EEG recordings and preprocessing* for detail).

### Behavioral Procedure

For each trial, participants were instructed to prepare two possible actions for upcoming tests by applying the single shared rule to two independent stimuli (Figure. 1a). The required two actions on each trial were selected from the shared action constellations or S-R mappings shown in Figure 1b.

**Fig. 1.**
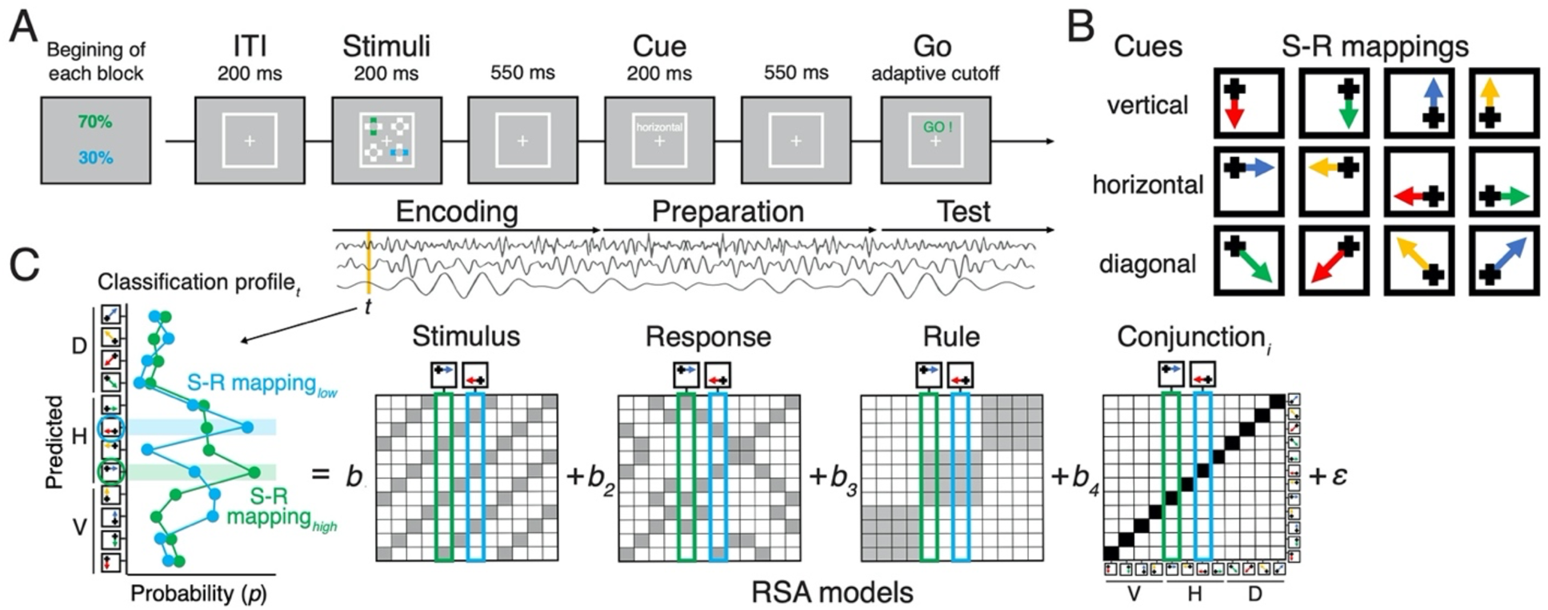
Task design and the procedure of decoding analysis. (A) Sequence of trial events in the rule-selection task with two independent action plans. Test probabilities of each action are assigned randomly every block. (B) Spatial translation of different rules (rows) mapping different stimuli (columns) to responses (arrows) (C) Schematic of the steps used for representational similarity analysis. For each sample time (*t*), a scalp-distributed pattern of EEG power was used to decode the specific rule/stimulus/response configuration of two actions of a given trial. The decoder produced sets of classification probabilities for each of the possible action constellations. The profile of classification probabilities reflects the similarity structure of the underlying representations, where action constellations with shared features are more likely to be confused. The figure shows an example of classification probabilities for two actions cued by a shared rule and two independent stimuli. For each trial and timepoint, the profile of classification probabilities was regressed onto model vectors as predictors that reflect the different, possible representations. In each model matrix, the shading of squares indicates the theoretically predicted classification probabilities (darker shading means higher probabilities) in all possible pairs of constellation. The coefficients associated with each predictor (i.e., *t-*values) reflect the unique variance explained by each of the constituent features and their conjunction for each action plan.

At the beginning of the *encoding* period (0 ms to 750 ms; Figure 1a), four crosses appeared at each of the four corners of a white square frame (6.6° in one side). Two of these locations were marked as to be remembered by coloring one of their cross bars either blue or green for 200ms, as shown in Figure 1a. Each bar was randomly assigned to be in a vertical or horizontal orientation, which allowed us to randomize and orthogonalize the locations of stimuli by including trials with an overlap of locations in 25% of trials. Then, following 550ms retention interval with only a fixation cross, one of the three possible action rules was randomly cued for 200ms followed by a 550ms delay. This marked the *preparation* period (750 ms to 1500 ms; Figure 1a) during which participants could prepare both actions based on combination of the rule and remembered locations. Each action rule (“vertical”, “horizontal”, and “diagonal”) uniquely mapped the four stimulus positions to four response keys that were arranged in 2 x 2 matrix (4, 5, 1, and 2 on the number pad; Figure 1b). For example, the horizontal rule mapped the left-top bar to the right-top response and the right-bottom bar to the left-bottom response as the correct set of responses. To focus on the effect of simultaneously planning two independent actions, we excluded trials during which two actions led to the same response due to the overlap of stimulus positions during analyses.

In order to encourage participants to prepare two actions before the final *test* period (starting 1500 ms after the stimulus, Figure 1a), an adaptive cut off was used to limit the test interval. The test screen showed a cue “GO!” in either blue or green to prompt participants to execute one of the prepared actions of the corresponding color. Responses were allowed to register only within a limited cutoff time interval following the cue. This cutoff interval was initially defined as 1200 ms, then it was adaptively adjusted trial-to-trial by decreasing the interval in any correct trials whereas decreasing the interval after five consecutive correct trials. The step size of adjustment of the interval was randomly selected from 11.8 ms, 23.5 ms, or 35.3 ms.

In order to manipulate priority, participants were explicitly instructed to prepare and maintain two actions simultaneously but to expect different probabilities of each action being tested. Expected test probability was cued by color of the bar stimuli (i.e., blue and green) to either a high (70%) or low (30%) test probability. The assignment of color to test probability was randomized for every block, therefore, one action plan was always more likely to be tested than the other in all trials. To prevent participants from disregarding information related to the low-test probability action, participants gained performance-based incentives only when the overall accuracy was above 85% in a given block, which was difficult to achieve if they used such a strategy, given the speeding required by the adaptive cutoff.

There were two practice blocks and 180 experimental blocks. Each block lasted 25 seconds, during which participants were instructed to complete as many trials as possible. Trials that were initiated within the 25 second block duration but extended beyond it were allowed to finish. Throughout the experimental session, participants were reminded to respond as accurately and fast as possible within adaptive cutoff intervals. Feedback about overall accuracy and the amount of performance-based incentives accrued was provided at the end of each block. All stimuli were generated in Matlab (Mathworks) using the Psychophysics Toolbox (Brainard, 1997) and were presented on a 17-inch CRT monitor (refresh rate: 60 *Hz*) at a viewing distance of 100 cm.

### EEG recordings and preprocessing

We recorded scalp EEG activities using 20 tin electrodes on an elastic cap (Electro-Caps) using the International 10/20 system. The 10/20 sites F3, Fz, F4, T3, C3, Cz, C4, T4, P3, PZ, P4, T5, T6, O1, and O2 were used along with five nonstandard sites: OL halfway between T5 and O1; OR halfway between T6 and O2; PO3 halfway between P3 and OL; PO4 halfway between P4 and OR; and POz halfway between PO3 and PO4. Electrodes placed ~1cm to the left and right of the external canthi of each eye recorded horizontal electrooculogram (EOG) to measure horizontal saccades. To detect blinks, vertical EOG was recorded from an electrode placed beneath the left eye. The left-mastoid was used as reference for all recording sites, and data were re-referenced off-line to the average of all scalp electrodes.

The scalp EEG and EOG were amplified with an SA Instrumentation amplifier with a bandpass of 0.01–80 *Hz*, and signals were digitized at 250 *Hz* in LabView 6.1 running on a PC. EEG data was first segmented into 27.5 second intervals to include all trials within a block. After time-frequency decomposition was performed, these epochs were further segmented into smaller epochs for each trial using the time interval of −200 ms to 2200 ms relative to the onset of stimuli (Figure 1a). The trial-to-trial epochs including blinks (>250 *μ*v, window size = 200 ms, window step = 50 ms), large eye movements (>1°, window size = 200 ms, window step = 10 ms), blocking of signals (range = −0.01 to 0.01 *μ*v, window size = 200 ms) were excluded from subsequent analyses.

### Time-Frequency Analysis

Temporal-spectral profiles of single-trial EEG data were computed via complex wavelet analysis (Cohen, 2014) by applying time-frequency analysis to preprocessed EEG data epoched for the entire block (>27.5 seconds to exclude the edge artifacts). The power spectrum was convolved with a series of complex Morlet wavelets), where *t* is time, *f* is frequency increased from 1 to 35 *Hz* in 35 logarithmically spaced steps, and σ defines the width of each frequency band, set according to *n*/2p*ft*, where *n* increased from 3 to 10. We used logarithmic scaling to keep the width across frequency band approximately equal, and the incremental number of wavelet cycles was used to balance temporal and frequency precision as a function of frequency of the wavelet. After convolution was performed in the frequency-domain, we took an inverse of the Fourier transform, resulting in complex signals in the time-domain. A frequency band-specific estimate at each time point was defined as the squared magnitude of the convolved signal for instantaneous power.

### Representational Similarity Analysis

We decoded action-relevant information in a time-resolved manner following our previously reported method with a few modifications (Kikumoto & Mayr, 2020). As the first step, separate linear decoders were trained to classify all possible action constellations (Figure 1b) for each independently planned action at every sample in trial-to-trial epochs. For each action, the 12 unique action constellations were defined by the combination of 3 rules and 4 stimulus positions (Figure 1b). This step produced a graded profile of classification probabilities for each action constellation, reflecting the multivariate distance of neural patterns between action plans. Decoders were trained with the instantaneous power of rhythmic EEG activity, which was averaged within the predefined ranges of frequency values (1-3 *Hz* for the delta-band, 4-7 *Hz* for the theta-band, 8-12 *Hz* for the alpha-band, 13-30 *Hz* for the beta-band, 31-35 *Hz* for the gamma-band), generating 100 features (5 frequency-bands X 20 electrodes) to learn. Within individuals and frequency-bands, the data points were *z*-transformed across electrodes to remove effects that uniformly influenced all electrodes. We used a k-fold repeated, cross-validation procedure to evaluate the decoding results (Mosteller & Tukey, 1968) by randomly partitioning single-trial EEG data into four independent folds with equal number of observations of each action constellation. All trials including error trials were used as the training sets. After all folds served as the test sets, each cross-validation cycle was repeated eight times, in which each step generated a new set of randomized folds. Resulting classification probabilities were averaged across all cross-validated results with the best-tuned hyperparameter to regularize the coefficients for the linear discriminant analysis.

We next performed representational similarity analysis (RSA) on the graded classification probabilities to assess the underlying similarity structure. Each RSA model matrix uniquely represents a potential, underlying representation (e.g., rules, stimuli, responses and conjunctions), which makes unique predictions for different action plans. Specifically, for independent decoding results of each of the action plans, we regressed the vector of logit-transformed classification probabilities onto RSA model vectors. To estimate the unique variance explained by competing models, we regressed all model vectors simultaneously, resulting in coefficients for each of the four model vectors. We also included subject-specific regressors of *z*-scored, average RTs and accuracies in each action constellation to reduce potential biases in decoding of conjunctions. These coefficients, expressed in their corresponding *t*-values, reflect the quality of action representations at the level of single trials, which was later related to variability in behavior. We excluded *t*-values that exceeded 5 SDs from means for each sample point, which excluded less than 1% of the entire samples. The resulting *t*-values were averaged over 40 ms non-overlapping time windows for visualization (Figure 3).

### Multilevel Modeling and Cluster-based Permutation Test

We used multilevel models to further assess the decoded representations (Figure 4; Table 1) and relate them to trial-to-trial variability in behavior (Figure 5). For these analyses, we used time-resolved and time-averaged RSA scores of both the basic action features and the conjunctions. For the results of time-resolved regression (Figure 4), we used non-parametric permutation tests to evaluate the decoding results in the time domain (Maris and Oostenveld, 2007). First, we performed a series of regression analysis over time and identified samples that exceeded threshold for cluster identification (cluster-forming threshold, *p* < .05). Then, empirical cluster-level statistics were obtained by taking the sum of *t*-values in each identified cluster with consecutive time points. Finally, nonparametric statistical tests were performed by computing a cluster-level *p*-value (cluster-significance threshold, *p* < .05, two-tailed) from the distributions of cluster-level statistics, which were obtained by Monte Carlo iterations of regression analysis with shuffled condition labels. When we regressed trial-to-trial response times and accuracies on RSA scores of action features, the coefficients were averaged across three time windows (as described in Behavioral Procedure) that were selected *a priori*: the encoding period, the preparation period, and the test period. These periods occurred 0 to 750 ms, 750 to 1500 ms, and 1500 to 2200 ms relative to the onset to the stimuli, respectively were used as predictors in the regression model. For all models, subject-specific intercept and slopes were included as random effects. For models predicting accuracies, we used multilevel logistic regression. For models predicting response times, error trials were excluded.

**Table 1.** Trial-by-trial RSA-scores of high and low-priority actions in each test probability context during the test phase.

| Decoded features | High-priority Tested |  | Low-priority Tested |  |
| --- | --- | --- | --- | --- |
|  | *b* (se) | *t*-value | *b* (se) | *t*-value |
| Rule | .071(.017) | 6.06*** | .062(.019) | 4.76** |
| Stimulus (high-priority) | .034(.010) | 3.41* | .021(.007) | 2.9* |
| Conjunction (high-priority) | .021(.006) | 3.57* | .007(.008) | 0.86 |
| Response (high-priority) | .034(.007) | 5.04** | .009(.009) | 1.04 |
| Stimulus (low-priority) | .011(.007) | 1.52 | .014(.006) | 2.19* |
| Conjunction (low-priority) | .003(.005) | 0.61 | .001(.007) | 0.125 |
| Response (low-priority) | .008(.005) | 1.61 | .015(.008) | 1.82* |

## Results

### Behavior

The pattern of behavioral results confirmed that the high priority action led to more efficient action selection (Figure 2). When the high priority action was tested as opposed to the low priority action, participants responded significantly faster, *F*(1,23) = 40.31, *MSE* = 962, *p* <.001., and produced fewer errors, *F*(1,23) = 31.81, *MSE* <.001, *p* <.001. Most of those errors were responses corresponding to the untested action plan rather than random responses. This suggests that participants encoded, and likely maintained, both action plans until the test period, but then occasionally selected the wrong plan. Such swapping errors occurred at significantly higher probability when the low priority action was tested, again consistent with participants prioritizing the high priority action during maintenance, *F*(1,23) = 12.59, *MSE* <.001, *p* =.002. Likewise, a trial-wise adaptive cutoff (i.e., response deadline) was significantly shorter for high priority action, *F*(1,23) = 33.72, *MSE* = 7479, *p* <.001. Therefore, the behavioral results indicate that both of the required actions were prepared, yet the action associated with the high test probability was prioritized over the other action.

**Fig. 2.**
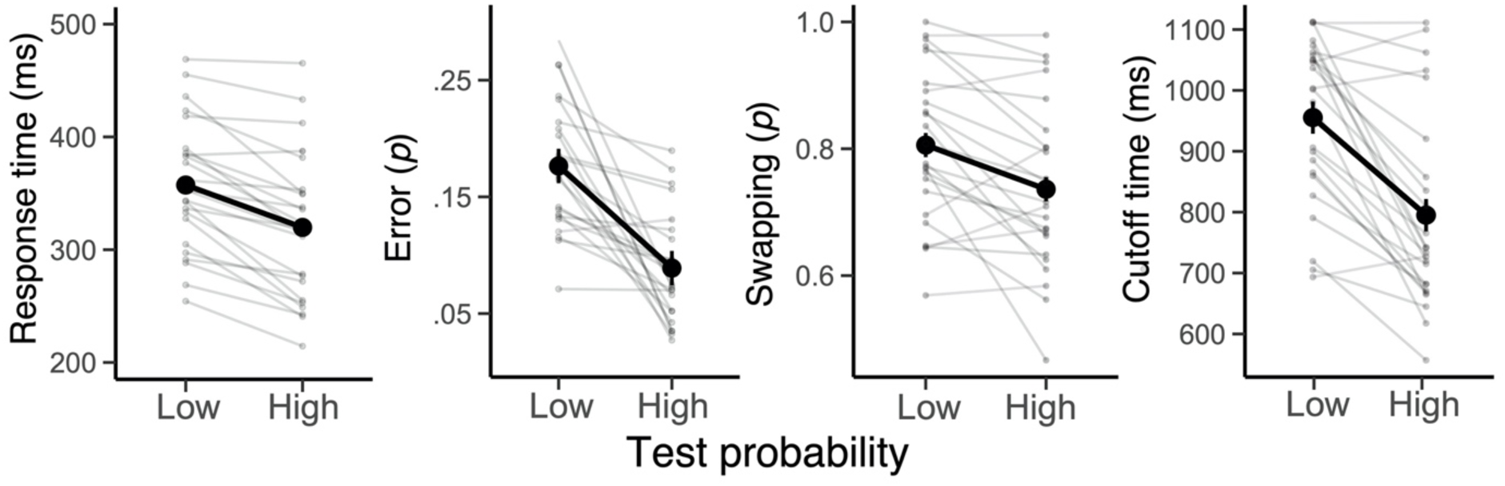
The effect of priority of actions on behavior. Average response times, error probability, swapping probability and cutoff time (i.e., trial-to-trial response deadlines) in the low and high test probability conditions. Note that a swapping error was computed out of all committed errors, rather than all trials. Error bars specify 95% within-subject confidence intervals. Faint lines plot individual participants.

### Action Representations during Early Encoding and Preparation Phase

To assess how two planned actions are maintained and selected via retro-cues, we decoded the representations of action features (i.e., stimuli, rules, responses and conjunctions for each action) from the time-resolved patterns of EEG activity using RSA. Then, we used a mixed-effect model to assess the effect of priority on the quality of representations and subsequent selection performance at the level of single trials.

The RSA revealed unique temporal trajectories for the individual features of high and low priority action plans (Figure 3). During the encoding phase, a mixed-effect model revealed that representations of two stimuli for each action were encoded, *t*(1,23) = 7.45, *b* = .183, 95% CI [.136 .230] for high priority; *t*(1,23) = 6.92, *b* = .168, 95% CI [.121 .216] for low priority; and they quickly decayed yet remained active during the preparation phase, *t*(1,23) = 3.65, *b* = .036, 95% CI [.016 .052] for high priority; *t*(1,23) = 3.20, *b* = .017, 95% CI [.006 .027] for low priority. During the preparation phase, the cued rule context shared by both plans was specified and maintained robustly, *t*(1,23) = 6.59, *b* = .144, 95% CI [.101 .186]. Further, distinguishable conjunctive representations for both required actions emerged, *t*(1,23) = 2.61, *b* = .142, 95% CI [.004 .025] for high priority; *t*(1,23) = 2.10, *b* = .008, 95% CI [.001 .016] for low priority. In contrast, the information about required responses was not reliably decodable during the preparation phase, *t*(1,23) = −.514, *b* = .002, 95% CI [−.010 .006] for high priority; *t*(1,23) = −1.57, *b* = −.008, 95% CI [−.017 .002] for low priority.

**Fig. 3.**
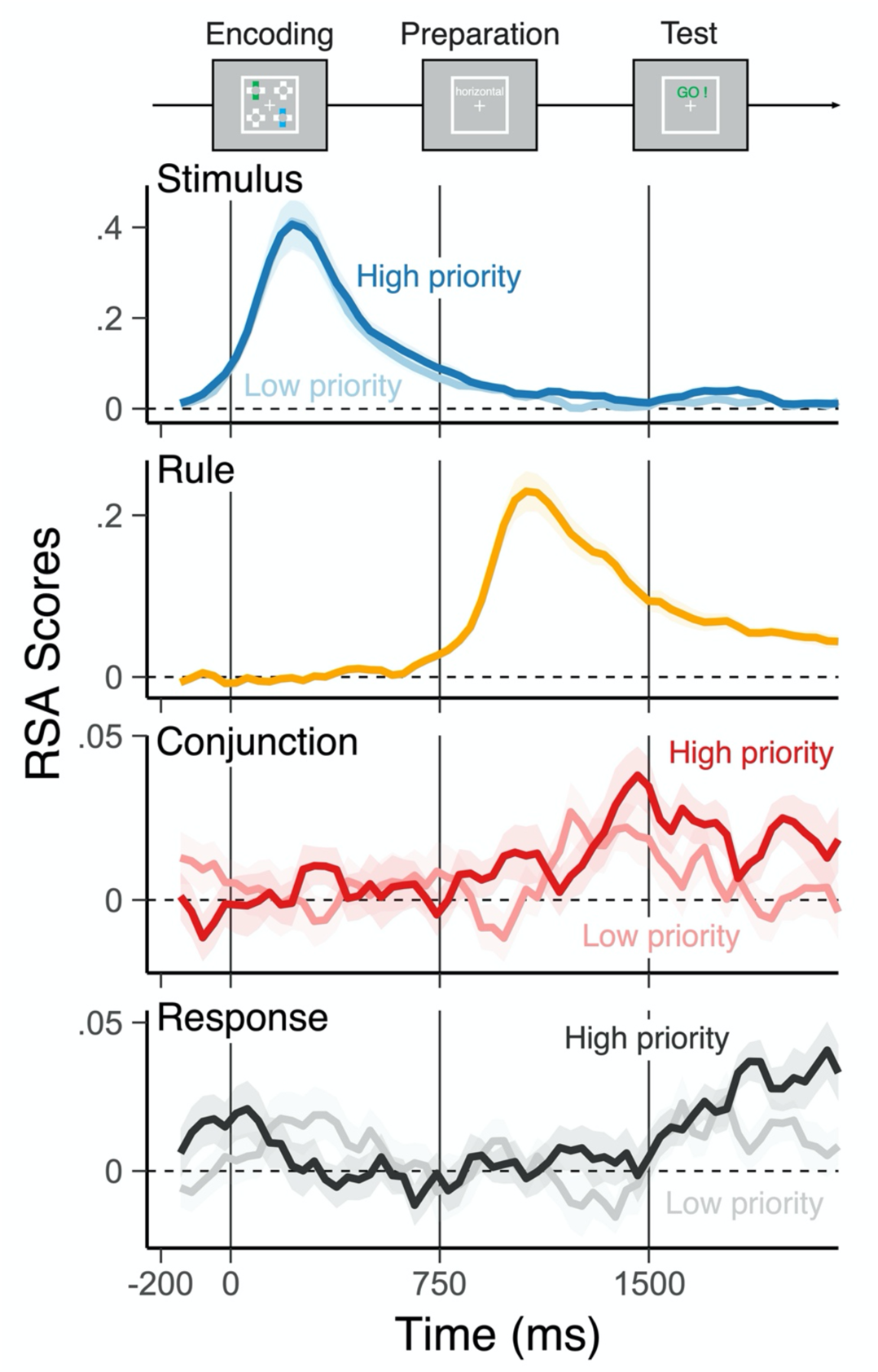
Trajectories of decoded representations of two actions in the different levels of priority. Average, single-trial *t*-values associated with each of the basic features (rule, stimulus, and response) and their conjunction derived from the RSA, separately for high priority (bold colors) action and low priority action (faint colors). Shaded regions specify the 95% within-subject confidence intervals.

Before the test phase, none of these representations were significantly modulated by different test probability, *t*(1,23) < 1.62, suggesting that neither action plan was selectively prioritized during preparation. Nonetheless, these results indicate early encoding and maintenance of conjunctive representations during the preparation phase and preceding the response. Note that in principle, candidate motor responses could have been fully prepared. Yet, action plans were maintained as conjunctive representations instead of response representations during the preparatory phase.

### Output gating of Action Representations During the Test Phase

In the test phase, participants were explicitly cued which prepared actions to select and made their response prior to the deadline (Figure 1A). We hypothesized that prioritization would modulate the states of selected and non-selected action representations. Specifically, we predicted that when a prepared conjunction is prioritized, this should facilitate selection of that plan, whereas a high-priority plan that is not selected should interfere with the execution of the one that is selected.

Although, on average, representations of individual features of high and low priority actions mostly became decodable only when they were cued as an output (Table 1), conjunctive representations were significantly modulated by the selection demand (i.e., cued as an output or not) dependent on their priority (Figure 4; see Table 2 for the main effects and Supplementary Figure 1 for the results using response-aligned data). Specifically, high priority items were more active than low priority items when they were cued, but the reverse was the case when low priority items were cued. Furthermore, we found that the strength of high and low priority conjunctions did not correlate with each other over trials across during the entire preparation and selection period, *t*(1,23) > −1.40. This result not only confirms that cross-over pattern of the results was not driven by direct competition among high and low priority conjunctions but also suggests that two conjunctions can be maintained in parallel and prioritization modulates output gating, not necessarily preparatory activation. A similar two-way interaction was observed for response representations but not for the stimulus representations (Figure 4). These results indicate that not every action feature is boosted by the retro-cue, and the conjunction is one of the targets of attentional modulation for selective output gating.

**Fig. 4.**
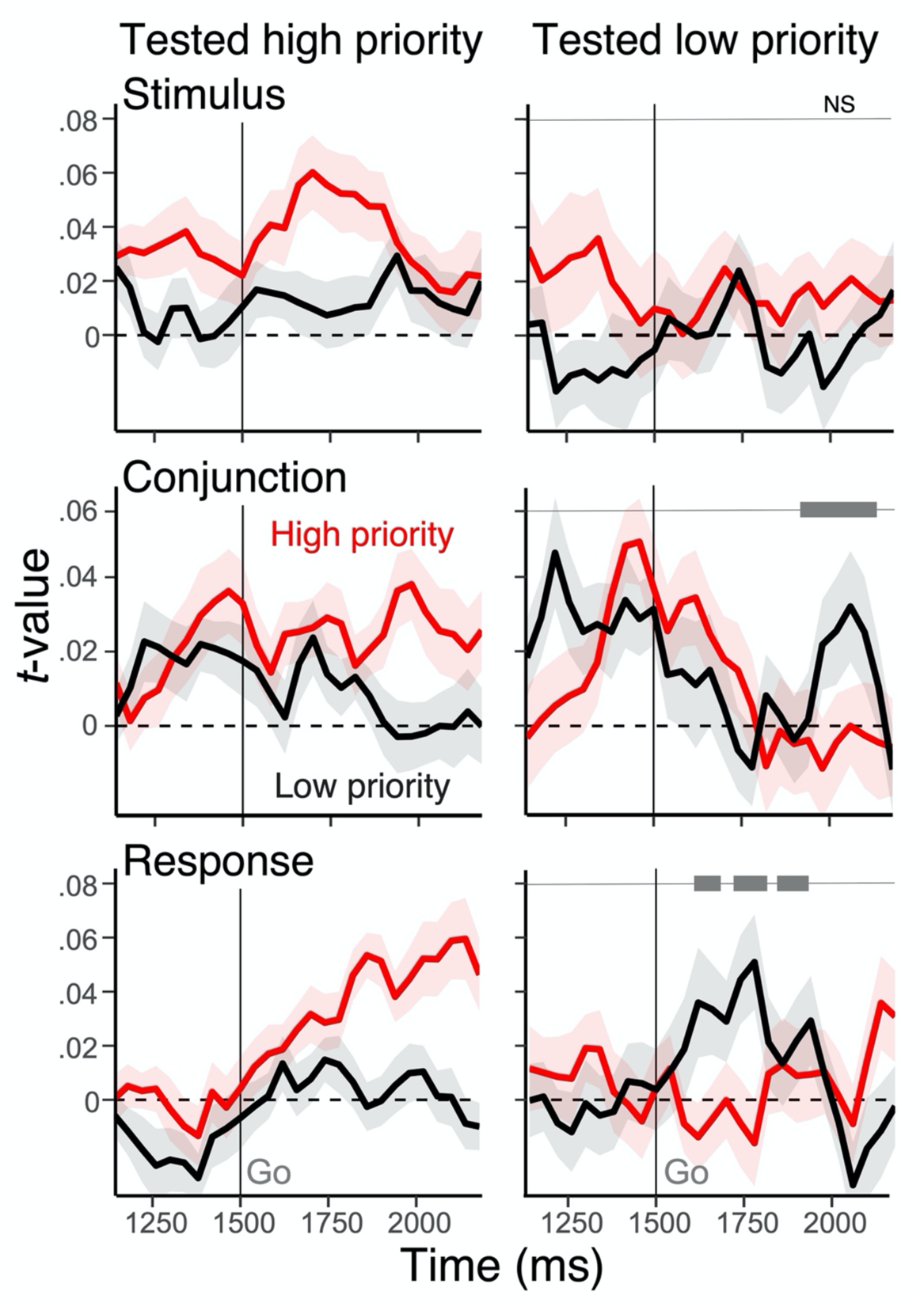
Modulation of action representations during selection. Average, single-trial *t*-values associated with the conjunction derived from the RSA, separately for high priority (red) action and low priority action (black). The left panels show RSA scores when the high priority action was tested, whereas the right panels show the result for the low priority action. Shaded regions specify the within-subject standard errors. On the right side panels, the gray bars at the top show clusters with a significant interaction effect between priority and the test type of actions after cluster-based correction (cluster *P*-value < .01).

**Table 2.** Cluster-level statistics for the main effects of the interaction model between test type and priority regressed on the RSA scores.

| Dependent variable | effect | time (ms) | cluster *T*-value | cluster *P*-value |
| --- | --- | --- | --- | --- |
| Stimulus | Priority | 1566 - 1598 | 20.11 | < .01 |
|  | Priority | 1846 - 1866 | 12.45 | < .01 |
| Conjunction | Test type | 1898 - 2050 | 98.36 | < .01 |
| Response | Priority | 1738 - 1794 | 38.70 | < .01 |
|  | Test type | 1858 - 1894 | 22.94 | < .01 |
|  | Test type | 2014 - 2110 | 67.02 | < .01 |

In addition, we observed an asymmetric interaction of action representations with different levels of priority on trial-to-trial response times and accuracies depending on the test type (Figure 5). Specifically, stronger representations of a stimulus, response and conjunction of the high priority action predicated faster and more accurate responses when that high priority action was tested by the cue. Such facilitatory effects were diminished or reversed into interference, when the low priority item was tested as a function of the strength of the high priority representation. In other words, on trials when the low priority action plan was cued, but the high priority action plan was strong, this led to slower responses and more errors.

**Fig. 5.**
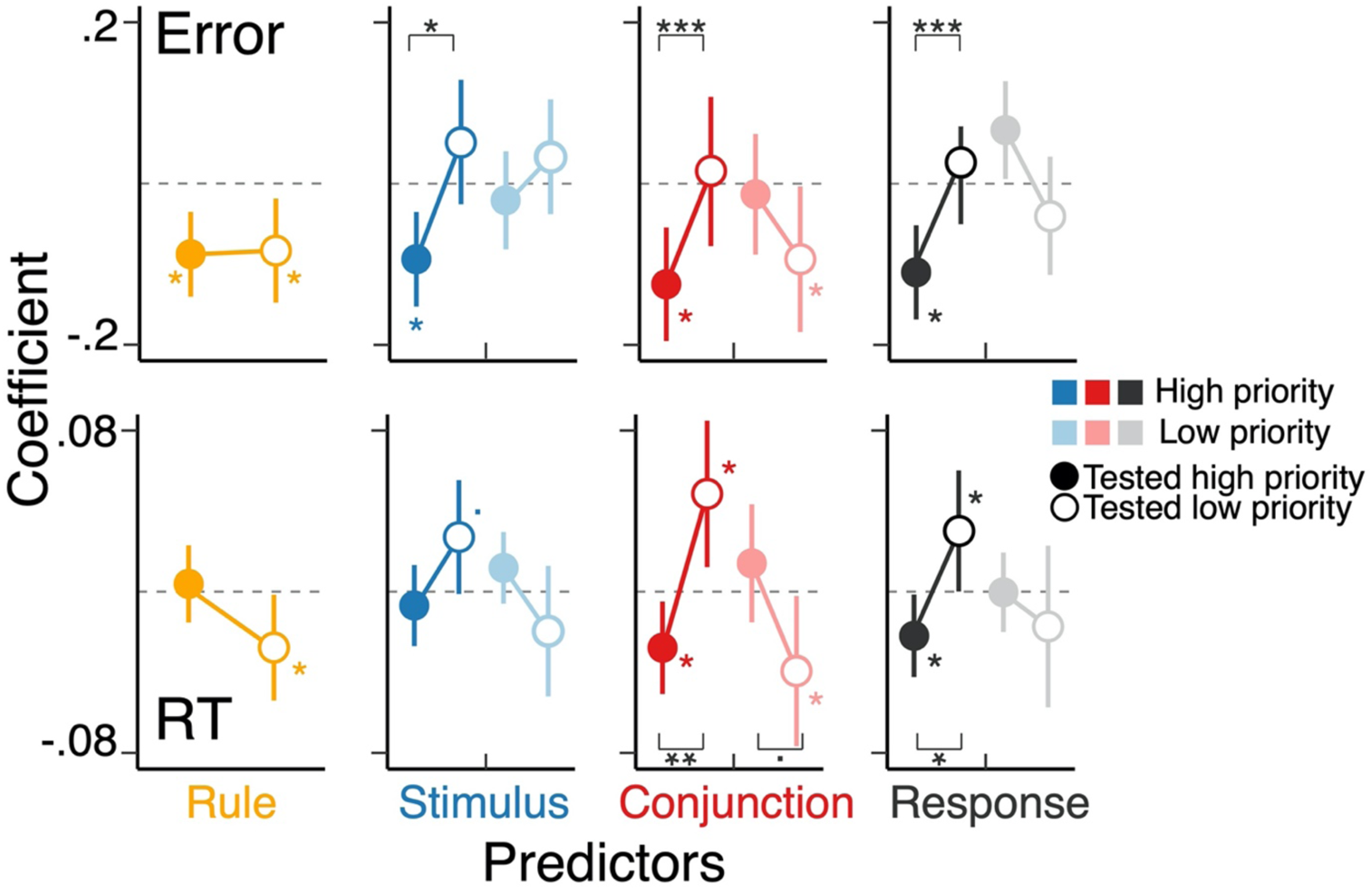
Interference between tested and untested actions with different level of priority. The coefficients of the multilevel regression models predicting the variability in trial-to-trial RTs and errors. The model included RSA scores of all action features of both actions during the test phase as predictors as well as the main effect of text context. The left-half side of a panel (denoted as “H”) for stimulus, conjunction, and response correspond to the features of high priority action, and the right-half side (denoted as “L”) shows the features of low priority action. The stars without a bracket indicate the level of significance for individual coefficients and the stars with a bracket shows the effect if selection (i.e., an action required to for the test).

We note again that when the high priority item was selected, the strength of all representations correlated with better behavior. In contrast, with the low priority item, only the strength of the conjunctive representation of the low priority item was predictive of behavior when that action was cued. There was no evidence of this relationship for the strength of the low priority response or stimulus representation. Furthermore, the strength of the representations of the low priority action mostly did not interfere with selection of the other, high priority action, except for the conjunction on RTs, although a quantitatively similar pattern was observed for the response representation.

Taken together, during selection for the output, the conjunction and response representations, but not stimulus representations, were selectively activated (Figure 4) and influenced efficient response selection (Figure 5) contingent on the priority and output gating demand of corresponding actions. These effects were asymmetric such that representations of high priority action tended to have a stronger impact on output selection, suggesting prioritization is established during output gating.

## Discussion

Our ability to prepare and select from a pool of action plans allows us to rapidly adapt to diverse and changing situations. Theories of cognitive control posit that an integrated control representation that incorporates a conjunction of task-relevant information, such as goals, action rules, sensorimotor features and rewards, may be critical for flexible action selection (Hommel 2019; Frings et al. 2020; Logan 1989). And, indeed, recent evidence has established the critical role of conjunctive representations in execution and control of responses (Kikumoto and Mayr 2020; Kikumoto and Mayr, 2022).

If conjunctive representations function as basic building blocks of action preparation and selection one would expect that people can maintain multiple conjunctions in working memory, prioritize them based on their expected utility, and selectively output one or another as circumstances dictate. The current study adapted a previously established paradigm for tracking conjunctive representations in order to test this hypothesis. We found evidence both for simultaneous preparation of two conjunctive representations and for priority-modulated output gating of the conjunction that corresponds with the currently targeted action. Furthermore, we found that conjunctive representations influence post-encoding output selection over and above the constituent representations, which were consistently targeted by attentional selection throughout a trial to maintain, prioritize and select an action from the candidates held in working memory.

Our results add to a growing body of evidence that maintenance of sensory information for upcoming output selection concurrently transform task-relevant sensory memories in a more action-oriented or proceduralized representational format (Oberauer 2009; Hommel 2019; Brass et al. 2017). Preparing future context-dependent actions recruited conjunctive representations that comprise the unique action constellation for output even after responses are fully specified (Figure 1, 4 and 5). Moreover, several current theoretical proposals for working memory suggest that encoded sensory information is reconfigured to a use-optimized format when a memorandum is assigned a specific use or selected for output (such as by retro-cueing) (Myers et al. 2017; Nobre and Stokes 2019; Orhan and Ma 2019). Consistent with these hypotheses, we observed here that participants can retrospectively prioritize and select conjunction representations based on task demands.

Notably, unlike prior studies that manipulate post-encoding selection from working memory (Griffin and Nobre 2003; Souza and Oberauer 2016; Ester et al. 2018), we observed that stimulus representations showed a relatively weak effect of priority and selection demand and a less of interference on behavior (Table 2; Figure 4; Figure 5). This may be partially driven by the fact that the sensory inputs were explicitly contextualized or transformed by the action rules for the task. However, another mutually compatible account of this discrepancy is that our RSA regression approach allows us to directly test conjunctive representations competing against other constituent formats, such as the simple stimulus representations.

Instead of other action features, our results showed conjunctive representations and response representations reflect the priority of planned actions during post-encoding selection. Consistent with recent theories that explained post-encoding priority effects by linking its informational content to associated actions through attention (Allport et al. 1987; Souza and Oberauer 2016; Olivers and Roelfsema 2020; González-García et al. 2020), these results suggest that more context-specific representations may be a better format to capture control processes for post-encoding selection of a planned action. Unlike typical memory tasks in which enhanced sensory representations suffice for successful completion of the task, realistic context-dependent actions require flexible control over input-output mapping. Beyond prioritization and selection, optimal control of actions may recruit conjunctive representations that minimally defines relevant task states to encode response representations that execute actions (Todorov 2004; McNamee and Wolpert 2019)

The importance of the integrated representation in our results also fit with recent theories about the geometry of task representations by nonlinear integration of information, wherein sensory and context signals are projected into a high-dimensional population code, as a key computational mechanism for working memory and cognitive flexibility (Fusi et al. 2016; Buschman 2021; Badre et al. 2021; Jazayeri and Ostojic 2021; Freund et al.). Supporting this view, neurons in association cortex (e.g., prefrontal and parietal cortex) exhibit nonlinear mixed-selectivity or coding of task feature information in a conjunctive manner. At the population level, this coding aids behavioral flexibility by discriminating readout for diverse combinations of inputs (Rigotti et al. 2013; Parthasarathy et al. 2019; Bernardi et al. 2020; Panichello and Buschman 2021).

A recent model of working memory explicitly addressed conjunctive coding by task-agnostic random networks and highlighted their role in flexibly maintaining arbitrary inputs (Bouchacourt and Buschman 2019). This model proposed that the flexible yet capacity-limited nature of working memory may originate from the interaction between a structured layer containing subnetworks of sensory features and an unstructured (random) layer responding to integrated information in high-dimensional spaces. Perhaps consistent with these perspectives, the observed conjunctive representations during preparation of future actions may reflect a working memory system that maximizes flexibility by relying on integrative connections (Buschman 2021).

A further benefit of high dimensional representations is that they can cast multiple task features into separable patterns, which could reduce their overlap and mitigate task interference (Musslick et al. 2020; Badre et al. 2021). In our task, the rule always overlapped between two action plans, thus action preparation could be aided by heightened separability in high dimensional representational geometry. Yet, we observed that when prioritized conjunctive representations were strong during the test phase, they interfere with selecting an alternate action (i.e., the low priority action), perhaps suggesting either that the separability between the two action plans was compromised or not complete. Though, it is also possible that the interference reflects downstream processes that must select between the action plans, as discussed below. Future studies should more directly investigate representational geometries encompassing two action plans and relate to the interference costs, including conditions that do not form integrated representations.

An important question left open by this work is what mechanism enacts prioritization and selection of these conjunctive representations. A context-dependent control mechanism that selectively gates into and out of working memory has been extensively studied as one such mechanism (O’Reilly and Frank 2006; Hazy et al.; Frank and Badre 2012; Badre and Frank 2012). One computational account, named the PBWM (prefrontal cortex basal ganglia working memory) model, implements gating of working memory via dopaminergic reinforcement learning in basal ganglia nested within cortico-striatal hierarchical loops. The computational architecture of the PBWM model provides a natural mechanism for combining values or expected utility of information into gating decisions. At the algorithmic level, the current study supports the use of selective output gating for action preparation. Yet, unlike previous studies focusing on the role of compositional representations of the task-relevant inputs (e.g., rule or context representations), our results highlighted an active role of integrated representations in biasing prioritization and selection of the planned actions on behavior.

Our results showed that multiple action plans could be retrospectively selected from within working memory, albeit with interference costs from the prioritized action during output selection phase. In the PBWM framework, the interference effect could be accounted for by changes in expected value that occur after the “GO !” cue modulating gating operations in the basal ganglia (e.g., changes in drift rate), which may occur regardless of the separability of cortical representations. The late appearance of the priority effect and its interaction with selection demand (Figure 4) and more pronounced interference effect of high priority action features (Figure 5) suggest that output selection of a low-priority action may induce switching of output gate that propagates down to the conflict at motor gating. From this perspective, the priority of actions is mainly reflected during gating operations for selecting one or another action. Consistent with this view, most errors were execution of the uncued actions (Figure 2). Though the present study cannot provide direct support for the involvement of biological gating circuits in the prioritization and selection of conjunctive representations, future studies could explicitly test this hypothesis using measures other than EEG and also could model the computational role of conjunctive representations in gating control of working memory.

Understanding cognitive flexibility requires answering how action plans are represented, maintained, prioritized, and selected based on changing circumstances in our environment. The current study provided an initial evidence that multiple future actions could be maintained as conjunctive representations, and this level of representation is a key target of processes that prioritize prepared actions and guide their retrospective selection.

## Supplementary Figures

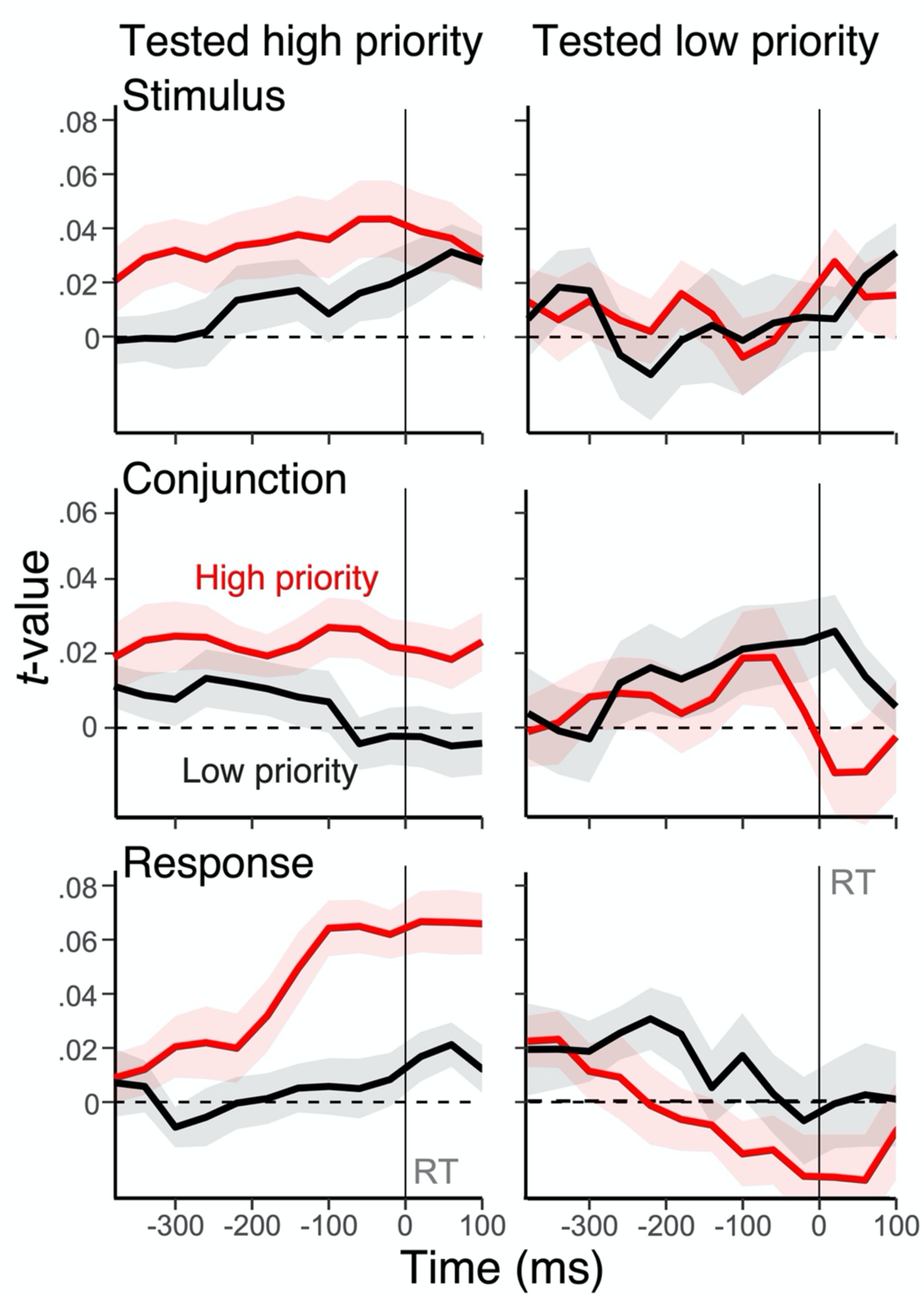
Modulation of action representations during selection response-aligned Figure 4 - figure supplement 1. Average, single-trial *t*-values associated with stimulus, response and conjunction derived from the RSA, separately for high priority (red) action and low priority action (black). The left panels show RSA scores when the high priority action was tested, whereas the right panels show the result for the low priority action. Shaded regions specify the within-subject standard errors.

## Acknowledgements

We would like to acknowledge the Cognitive Dynamics lab and the Badre Lab, particularly Jiafan Jia, Apoorva Bhandari, Haley Keglovits, for helpful comments and discussion. This project was supported by funding from the National Institute of Mental Health (R01 MH125497), the National Institute of Neurological Disorders and Stroke (R21 NS108380), and a Multidisciplinary University Research Initiative award from the Office of Naval Research (N00014-16-1-2832) to DB and from the National Science Foundation Grant(1734264) to UM.

